# Pangenomic read mapping

**DOI:** 10.1101/813634

**Authors:** Siavash Sheikhizadeh Anari, Dick de Ridder, M. Eric Schranz, Sandra Smit

## Abstract

In modern genomics, mapping reads to a single reference genome is common practice. However, a reference genome does not necessarily accurately represent a population or species and as a result a substantial percentage of reads often cannot be mapped. A number of graph-based variation-aware mapping methods have recently been proposed to remedy this. Here, we propose an alternative multi-reference approach, which aligns reads to large collections of genomes simultaneously. Our approach, an extension to our pangenomics suite PanTools (https://git.wur.nl/bioinformatics/pantools), is as accurate as state-of the-art tools but more efficient on large numbers of genomes. We successfully applied PanTools to map genomic and metagenomic reads to large collections of viral, archaeal, bacterial, fungal and plant genomes.

## Background

Mapping short reads against a reference genome is the starting point of almost all quantitative and comparative genomics pipelines (1). However, it suffers from a systematic bias towards the reference alleles (often referred to as reference bias): reads which are highly polymorphic or totally absent in the reference are discarded. This means variants that could be of great value, for example in disease diagnostics in humans or resistance genes in plants, are potentially overlooked. Mapping reads against multiple genomes representing a genus, a species or a population, i.e. a pangenome, partially addresses this problem, allowing the detection of variants that would not be detected using a single reference. A variety of graph-based pangenomic approaches have therefore emerged recently, which can generally be categorized as either variation-aware or multi-reference. Variation-aware mapping is useful for applications in which variation between individuals is limited and measured extensively, such as in human genetics (2). Multi-reference approaches in contrast are more suited for applications in which more divergent individuals are studied, such as in comparative genomics (3).

BWBBLE (4) is a variation-aware method that maps reads against a BWT-indexed linear multi-genome reference, built from one reference and a set of variant (VCF) files. Graphtyper (5) iteratively enriches a variation-aware graph with known or discovered variants for read mapping and population-scale genotyping. Similarly, the variation graph toolkit (6) constructs a bi-directed variation-aware graph as a reference to improve the accuracy of mapping, specifically in highly polymorphic regions. GenomeMapper (7) is the first multi-reference approach that represents a reference genome and its differences to a set of other genomes in a hash-based graph structure against which reads can be aligned. GCSA (8) converts a multiple sequence alignment (MSA) of genomes into a finite automaton which is BWT-indexed to allow pattern search. PanVC (9) uses the MSA as a pangenome reference and map reads against the matrix, where the heaviest path serves as an *ad hoc* reference to improve the accuracy of downstream variant callers.

These approaches, mostly targeting the human genome and/or focusing on specific variable regions, demonstrate that using broader reference representations can improve read mapping. However, they do not suffice to study collections of individual genomes of highly dynamic species such as fungi and plants. In such collections, genome co-linearity is often not preserved; moreover, scalability becomes an issue as the number of genomes grows. To tackle these issues, we propose a multi-reference read mapping approach, as an extension to PanTools. Briefly, PanTools is a suite of tools for large-scale comparative analysis building on a pangenome representation stored in a graph database (10). Our mapping method can align millions of short reads to hundreds of eukaryotic or thousands of prokaryotic genomes simultaneously, producing one SAM/BAM file per genome. It also provides a competitive mapping mode, which is useful for abundance estimation and binning in metagenomics samples. We demonstrate that PanTools is as accurate as the state-of the-art read mappers and per-genome mapping time decreases with increasing numbers of genomes.

## Results

We have extended our pangenome tool suite, PanTools, with a method to efficiently map genomic reads against multiple genomes in a graph-based representation (the algorithm is described under Methods). Conceptually, this eliminates the strong reference bias, which stems from mapping to a single genome. Reads that do not map on one genome may map on another genome, yielding a more complete picture of the genomic makeup of a sample. In application, PanTools offers two advantages that allow mapping efficiency to improve as the number of genomes grows. First, a single joint *k*-mer index is available for all genomes, resulting in fast identification of candidate hits that can then be targeted for full alignment. Second, redundant sequence alignments are avoided by recording previous alignments; if two genomes are similar, fewer alignments have to be made.

PanTools features two modes of read mapping. In ‘normal’ mode, genomic reads are independently mapped against all genomes in the pangenome, identifying the most likely mapping location of each read in each genome. In contrast, in ‘competitive’ mode reads are mapped to the most likely location in the entire pangenome, such that a read with the highest mapping score on genome A will not be mapped to genome B with a lower score. Competitive mapping is useful in various applications involving mixed samples, such as metagenomics samples, pathogen/host samples, or nuclear/organellar samples.

Here we present the performance of PanTools as a multi-genome read mapper on various sets of simulated and real data from bacteria, fungi and plants. We describe the accuracy and runtime of our approach, compared to a number of other read mappers. In addition, we present two use cases, demonstrating scalability to large genomes and application in metagenomics.

### PanTools is as effective as current read mappers

To learn about accuracy and speed of pangenomic read mapping compared to state-of-the-art single-reference mappers, we simulated read data from two Illumina platforms (HiSeq 2500 and MiSeq v3) and mapped these against the reference genomes of *E. coli* and *S. cerevisiae*. There is often a trade-off between runtime and accuracy of read-mappers, i.e. more accurate results can be attained by using more sensitive settings at the cost of a higher runtime.

This experiment demonstrates that, on single genomes, PanTools is competitive with widely used methods. Figure 1 shows time-accuracy plots of five read mappers (running with default settings on a single processing core): PanTools; two popular BWT-based methods, BWA-MEM (11) and Bowtie2 (12); and two hash-based mappers, Stampy (13) and NextGenMap (14). Stampy was the slowest method when applied to HiSeq data, followed by Bowtie2 which was over two times faster, yet approximately four times slower than the remaining methods. On the MiSeq data, Bowtie2 was slowest, but the overall picture was similar. In terms of accuracy, almost all methods perform similarly with the exception of Bowtie2 on MiSeq data. PanTools, BWA-MEM and NextGenMap are highly comparable in terms of speed and accuracy (Additional file 1: Experiment 1).

**Figure 1:**
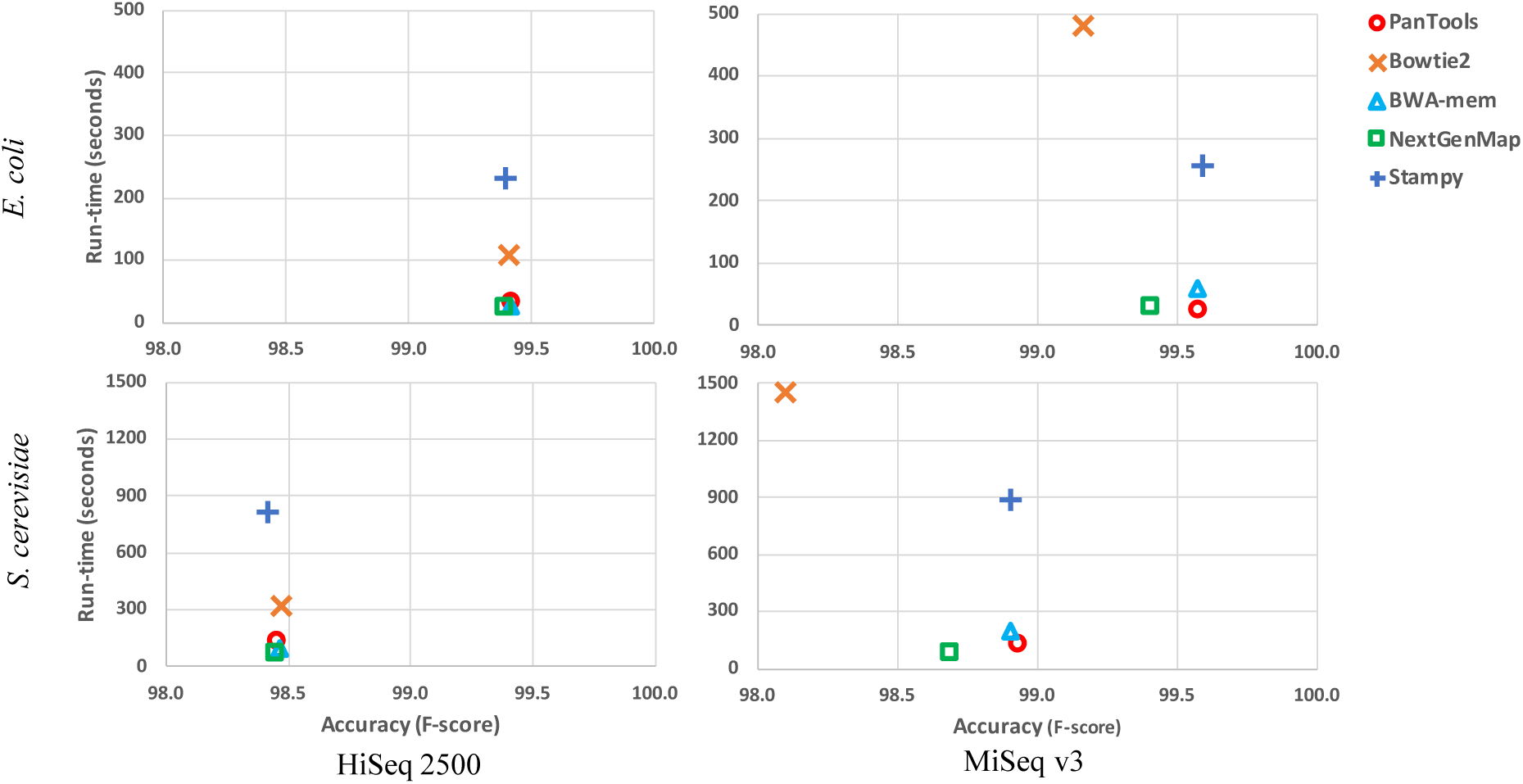
Runtime versus accuracy of five read mappers on four simulated Illumina datasets. Accuracy is presented in terms of the F-score (see Methods).

### The pangenomic approach becomes more efficient as the number of genomes grows

We compared the scalability of the best performing tools (PanTools, BWA-MEM and NextGenMap) to large sets of genomes. To learn about the effect of evolutionary distance between genomes, we mapped simulated *S. cerevisiae* reads against four pangenomes of ten fungi chosen at the levels of strain (ST), species (SP), genus (GN) and family (FM) (see Additional file 1: Experiment 2). Figure 2A shows the average runtime per genome of mapping reads against 1-10 genomes. For PanTools, as a multi-genome read mapper, this time is calculated as the total runtime divided by the number of genomes in each experiment. For the singe-genome read mappers, BWA-MEM and NextGenMap, it reflects the average of runtimes up until each point. All tools were running with 8 threads.

**Figure 2:**
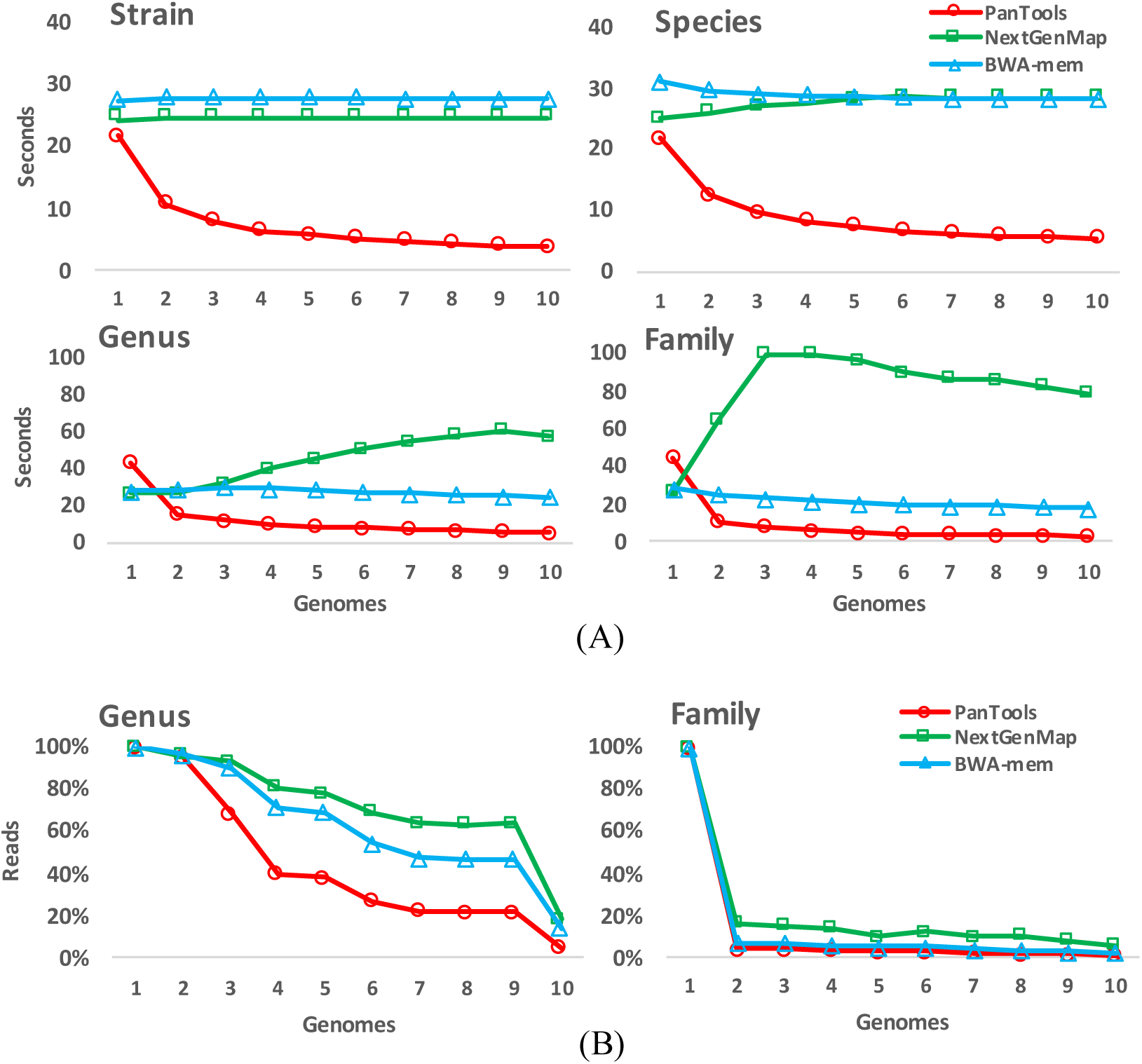
Mapping simulated reads from *S. cerevisiae* strain S288C on four pangenomes of ten fungi. **A.** Runtime of three methods at the level of strain S288C, species *S. cerevisiae*, genus *Saccharomyces*, and family Saccharomycetaceae. Genomes are sorted in decreasing order by the average number of reads that the tools managed to map, roughly reflecting their similarity to the reference genome of *S. cerevisiae*. **B.** The mapping percentage depends on the similarity between the sequenced strain and the reference genome and on the default sensitivity of the methods.

In PanTools the runtime per genome decreased when the number of genomes in the pangenome increased; the more related the genomes were, the higher the speedup. The runtime of BWA-MEM was very stable, around 30 seconds per genome, whereas that of NextGenMap radically increased as more divergent genomes were included in the set. All the tools had very similar mapping percentages at the strain and species levels, yet NexGenMap had the highest mapping percentage on diverged genomes at the genus and family levels (Figure 2B), correlated with its high runtime. The mapping percentage of PanTools can likewise be increased (at the cost of a higher runtime) through parameter settings. However, we chose less sensitive default settings, because in many applications read mapping is limited to the species level. PanTools parameters and settings are described in detail in the Methods.

PanTools’ speed is largely due to avoiding redundant alignments by maintaining a list of previous alignments. When the constituent genomes are closely related, the chance of finding an alignment in this list is high. Table 1 shows that this approach saves computations in this experiment, in particular when genomes are highly similar.

**Table 1:**
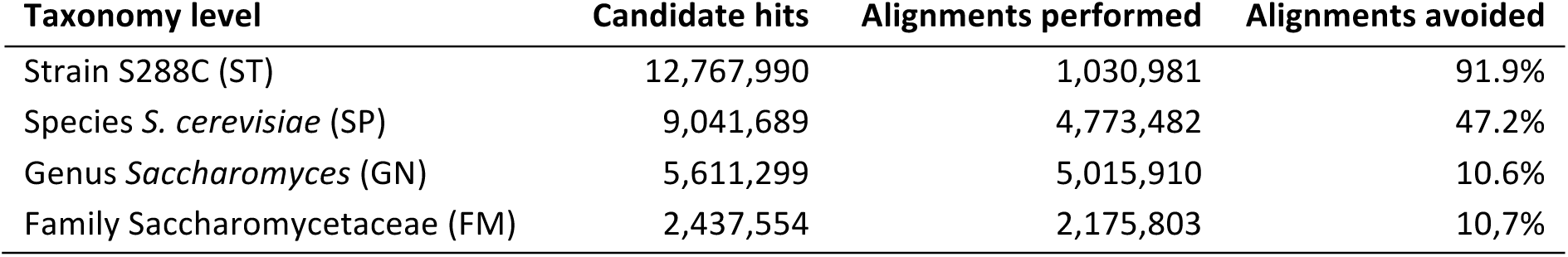
PanTools avoids redundant sequence alignments.

### Use case 1: Pangenomic read mapping in plants

To illustrate the utility of mapping to a panel of genomes rather than to a single reference, we applied PanTools to a case typically encountered in plant genomics: mapping reads of various (often relatively distant) accessions to a reference genome. We started with the model plant, *Arabidopsis thaliana*, and mapped reads of accession DJA-1 (Illumina library ERR2721960) to the reference genome Col-0. In this experiment, 12.28% of reads could not be mapped, potentially preventing the discovery of important variants. We next mapped these unmapped reads to the pangenome of 19 accessions of *A. thaliana* (15) and managed to map ∼3% of these. Finally, we clustered the overlapping reads mapped to each accession to detect moderately covered genomic regions (coverage ≥ 10), which are either absent from or highly different in the reference Col-0. We detected on average 844 such regions with an average size 389bp in each of the other accessions (Table 2).

**Table 2:** Reads unmapped to the *A. thaliana* reference Col-0 re-aligned to a 19 accessions pangenome.

| Genome | Accession | Number of reads | Mapping percentage | Number of regions | Length of regions (min-avg-max) |
| --- | --- | --- | --- | --- | --- |
| 1 | Col-0 | - | - | - | - |
| 2 | Bur-0 | 112,710 | 3.44 | 951 | (151-406-2917) |
| 3 | Can-0 | 121,623 | 3.72 | 1,048 | (151-377-2917) |
| 4 | Ct-1 | 87,082 | 2.66 | 603 | (151-415-2917) |
| 5 | Edi-0 | 109,819 | 3.36 | 957 | (151-397-6124) |
| 6 | Hi-0 | 79,741 | 2.44 | 608 | (151-370-2918) |
| 7 | Kn-0 | 108,922 | 3.33 | 918 | (151-375-2918) |
| 8 | Ler-0 | 110,729 | 3.38 | 929 | (151-378-5000) |
| 9 | Mt-0 | 92,108 | 2.81 | 715 | (151-349-2574) |
| 10 | No-0 | 98,889 | 3.02 | 845 | (151-404-3306) |
| 11 | Oy-0 | 85,031 | 2.60 | 610 | (151-398-3103) |
| 12 | Po-0 | 84,054 | 2.57 | 698 | (151-365-2085) |
| 13 | Rsch-4 | 97,224 | 2.97 | 817 | (151-406-2917) |
| 14 | Sf-2 | 117,301 | 3.58 | 1063 | (151-373-2897) |
| 15 | Tsu-0 | 106,901 | 3.27 | 924 | (151-392-2903) |
| 16 | Wil-2 | 113,349 | 3.46 | 950 | (151-379-2903) |
| 17 | Ws-0 | 112,773 | 3.45 | 888 | (151-403-2917) |
| 18 | Wu-0 | 100,646 | 3.08 | 782 | (151-409-2917) |
| 19 | Zu-0 | 108,042 | 3.30 | 882 | (151-400-2917) |

As an example, we found a region of 248 base pairs absent in the reference Col-0 (first row, position Chr1:24,201,231) as well as in accession Wil-2 (position Chr1:23,713,822), but present in all other accessions. Figure 3 shows the multiple sequence alignment of this region in all accessions and reads which failed to be mapped against this region in Col-0 and Wil-2 but mapped to the other accessions. Similarly, Figure 4 shows such an alignment for a region of length 283bp in the sequenced DJA-1 individual, which is highly mutated in other accessions, specifically in Col-0. None of the 36 reads covering this region were mapped to Col-0, however between 24-33 reads were mapped to the other 18 accessions. There were 24 SNPs and 7 short indels in the alignment of the assembled reads and the corresponding region in Col-0, which explains the read mapping problems.

**Figure 3:**
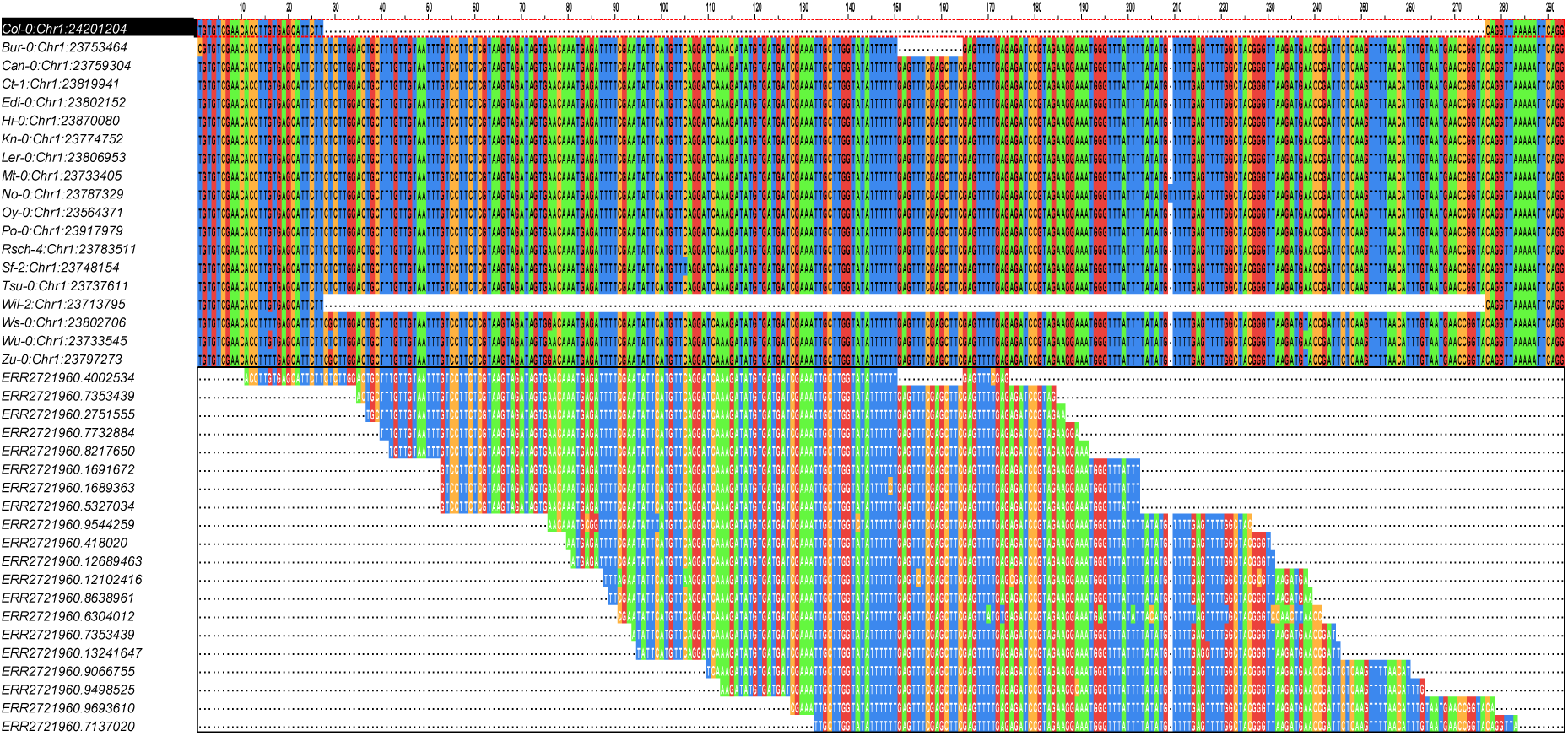
A deletion of 248 bp in chromosome 1 of the reference Col-0 and accession Wil-2.

**Figure 4:**
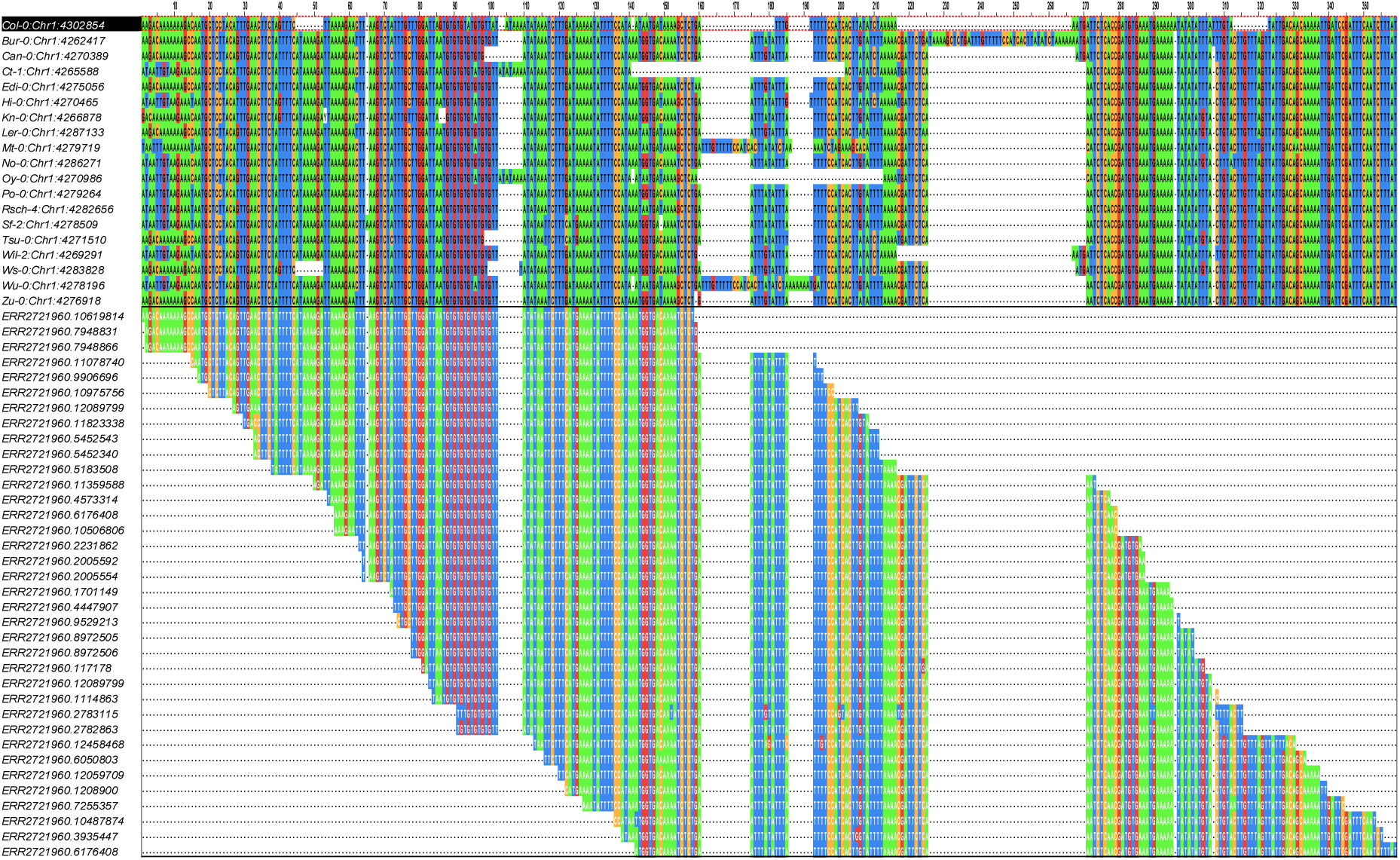
A highly mutated region starting at Chr1:4,302,854 in the reference Col-0. The large number of SNPs and indels in Col-0 prohibits alignment of reads that can be mostly mapped to the other accessions.

Next, we investigated whether PanTools’ read mapping would scale to larger, more complex genomes. To this end, we mapped three large paired-end sequencing libraries from the 150 Tomato Genome Resequencing Project (3) to the reference genome of tomato *Solanum lycopersicum* (Heinz 1706) and the three additional species *Solanum pennellii* (LA716), *Solanum pimpinellifolium* (LA480), and *Solanum habrochaites* (LYC4). PanTools achieved a high mapping percentage and additionally captured a large number of regions absent in the reference, or present but highly variable (Table 3).

**Table 3:**
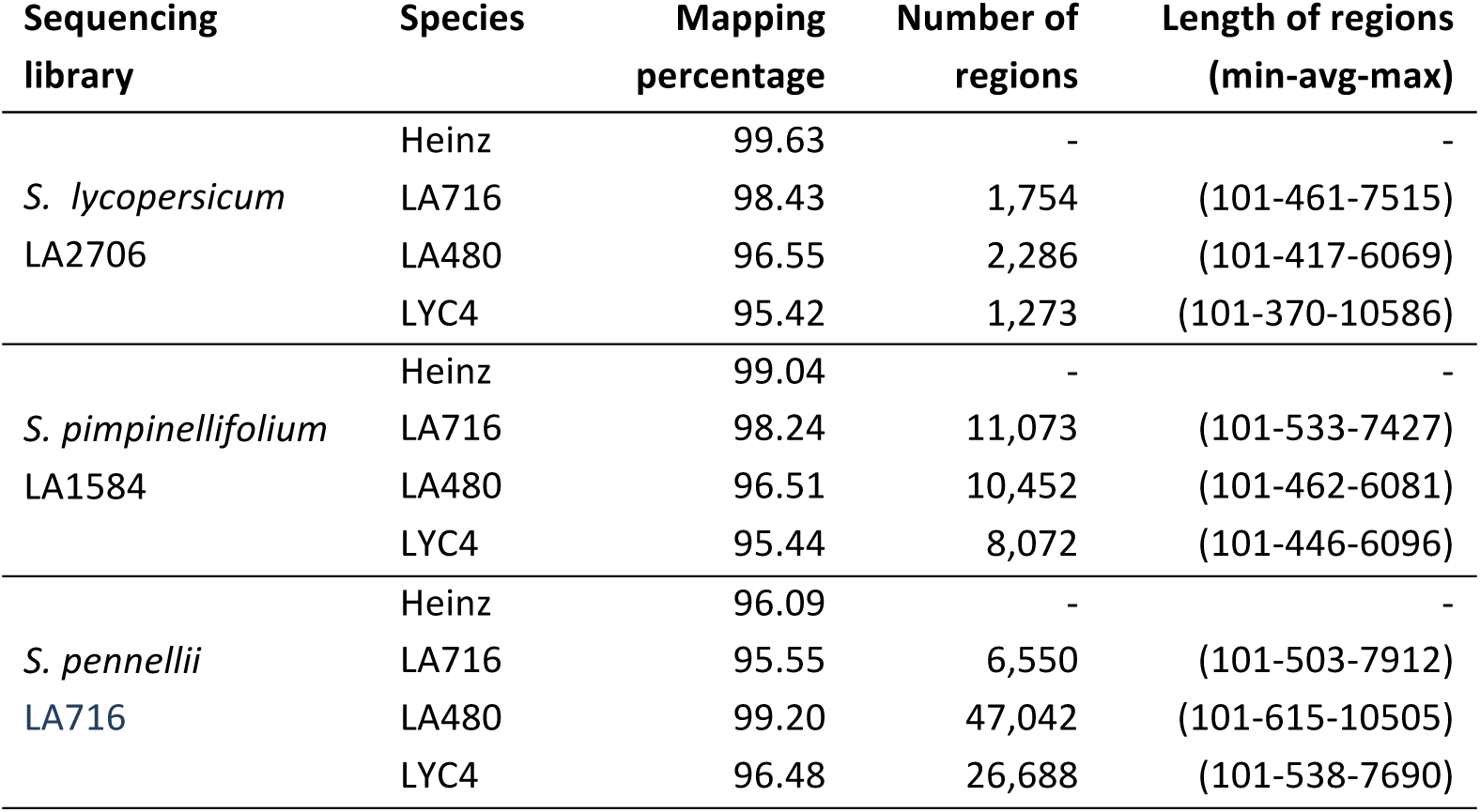
Mapping three libraries to a pangenome of four tomato accessions. The number and length of detected absent/highly variable regions in each reference are given in the last two columns.

| Sequencing library | Species | Mapping percentage | Number of regions | Length of regions (min-avg-max) |
| --- | --- | --- | --- | --- |
| *S. lycopersicum*<br>LA2706 | Heinz | 99.63 | - | - |
|  | LA716 | 98.43 | 1,754 | (101-461-7515) |
|  | LA480 | 96.55 | 2,286 | (101-417-6069) |
|  | LYC4 | 95.42 | 1,273 | (101-370-10586) |
| *S. pimpinellifolium*<br>LA1584 | Heinz | 99.04 | - | - |
|  | LA716 | 98.24 | 11,073 | (101-533-7427) |
|  | LA480 | 96.51 | 10,452 | (101-462-6081) |
|  | LYC4 | 95.44 | 8,072 | (101-446-6096) |
| *S. pennellii*<br>LA716 | Heinz | 96.09 | - | - |
|  | LA716 | 95.55 | 6,550 | (101-503-7912) |
|  | LA480 | 99.20 | 47,042 | (101-615-10505) |
|  | LYC4 | 96.48 | 26,688 | (101-538-7690) |

### Use case 2: Abundance estimation and binning of metagenomics data

In metagenomics studies the goal is often to identify the constituent organisms at a specific taxonomic level (e.g. by binning) and estimate their abundances. Numerous pipelines are available, usually based on targeted sequencing of the 16S ribosomal gene or on whole metagenome shotgun (WMGS) sequencing (16). In the latter case, mapping to a set of reference genomes is an extremely computationally intensive step. PanTools provides a competitive mapping mode to support such analyses. To demonstrate its use, we competitively mapped a metagenomics stool sample (SRS011061) from HMRARG2 (17) on the pangenome of the reference genome database of the Human Microbiome Project (HMP)b (see Methods). A list of all strains and their estimated abundances is available in Additional file 1: Experiment 5. We found a strong correlation between our estimated abundances and those found in the HMSCP report (19) (Supplementary Figure 1). Two bacterial strains, *Parabacteroides merdae* (ATCC 43184) and *Bacteroides cellulosilyticus* (DSM 14838), were the most abundant strains in this sample.

We also evaluated the accuracy of abundance estimates of PanTools on three benchmark data sets provided by the CAMI (Critical Assessment of Metagenome Interpretation) initiative (20), of low, medium and high complexity and compared it to those of two tools specifically developed for this problem, Kallisto (21) and DiTASiC (22). We ran PanTools in two *random-best* competitive modes; in the first run, we uniformly distributed shared reads between genomes, where in the second we considered the coverage of uniquely mapped reads in the first run to calculate the probabilities by which shared reads are assigned to the genomes. The idea behind this approach was that unique reads come from the strain-specific regions of the genomes and their abundance reflects the relative abundance of the genome in the sample. This approach significantly improved the accuracy of PanTools on the medium complexity CAMI data set, where there was a large imbalance between the abundance of some extremely similar strains.

Table 4 shows the accuracy of the abundance estimates of Kallisto, DiTASiC and PanTools (see Supplementary Figure 2) in terms of the root mean squared error and correlation coefficient between estimates and the ground truth. In this experiment, Kallisto performed best in terms of speed and accuracy. The runtime of DiTASiC grows quadratically with the number of genomes, as it needs to calculate the pairwise similarity between the genomes. On the high-complexity dataset of 1074 genomes we killed the process after 10 days. In contrast, PanTools was able to handle all the three datasets in reasonable time with accuracy comparable to Kallisto, while additionally simultaneously binning reads in individual SAM files which could in principle directly be passed on to an assembler to build the contigs.

**Table 4:** Root mean square error (RMSE) and correlation coefficient between estimates and ground truth and runtime of three methods on the three CAMI benchmark datasets.

| Dataset complexity | Tool | RMSE | Correlation coefficient | Runtime (seconds) |
| --- | --- | --- | --- | --- |
| **Low** | Kallisto | 121,072 | **0.999** | 361 |
|  | DiTASiC | 205,980 | 0.998 | 2,820 |
|  | PanTools | 367,277 | 0.994 | 2,263 |
|  | PanTools (coverage-based) | 148,138 | **0.999** | 4,619 |
| **Medium** | Kallisto | 141,309 | 0.996 | 542 |
|  | DiTASiC | 121,824 | **0.997** | 52,200 |
|  | PanTools | 707,460 | 0.881 | 2,218 |
|  | PanTools (coverage-based) | 375,896 | 0.969 | 4,479 |
| **High** | Kallisto | 35,960 | **0.980** | 1,321 |
|  | DiTASiC | - | - | > 10 days |
|  | PanTools | 42,404 | 0.972 | 5,585 |
|  | PanTools (coverage-based) | 44,866 | 0.969 | 11,112 |

## Discussion

Multi-genome read mapping is necessary to overcome the “reference bias” that comes from only considering reads that map to a single reference. Unmapped reads are typically not considered for downstream analyses, while these could point to interesting variants. As we have demonstrated, unmapped reads in a sample can originate from genomic regions absent in or highly different from the reference. Ideally, the known variation between different genomes is exploited to improve read mapping across these regions. Existing variation-aware read mappers, such as Graphtyper (5), enrich a reference genome with known variants to improve read mapping across highly variable regions and capture polymorphisms, which are finally called with respect to the reference. This approach works well for specific genomic regions, e.g. HLA genes in human, where many variants are already known (23).

Still, reads from regions not present in such enriched references will remain unmapped. In studies on species with highly dynamic genomes, e.g. crops, where gene content varies and co-linearity is typically not preserved, a multi-genome read mapping approach is therefore preferable. PanTools is not variation-aware in the sense that information from genome A is used to map a read to genome B, but it efficiently maps reads to all genomes in a (potentially large) set. A set of reads may be mapped to a region in genome B, while they do not map on the homologous region in genome A because of SNPs and indels (as shown in Figure 4). By detecting homologous regions between genomes, it is, in principle, possible to project reads from one genome to the corresponding region in another genome, resembling the results of variation-aware methods.

PanTools can simultaneously generate alignment files (SAM/BAM) for multiple genomes. These can be fed to any variant caller to detect variants with respect to all the constituent genomes. Variants not captured in one reference thus may be found with respect to one or more of the other genomes. PanTools scales well to thousands of complete bacterial or fungal genomes and to collections of large genomes, such as those of plants. However, interacting with extremely large databases (e.g. tens of plant genomes) is time-consuming, as the database cannot be fully buffered in memory. A high repeat content of genomes also increases the runtime, as it causes certain nodes to have many genomic locations. In our experiments with the pangenome of four tomato accessions, we overcame this limitation by ignoring low-complexity nodes when collecting candidate hits.

Our current method is designed to map genomic short reads, single or paired-end. Soft clipping has been implemented, but split alignments are not reported. We intend to develop this further in the future, as it is required for the detection of (some forms of) structural variation. Along the same lines, we will investigate spliced mapping of transcriptome data, considering multiple partial hits per read. Mapping long reads, e.g. PacBio or Oxford Nanopore, would be another useful extension, but this requires additional work, for example to implement an additional *k*-mer index with smaller *k* to handle higher rates of error and an alternative (banded) alignment approach which would scale to longer sequences.

## Conclusions

The number of sequenced species is increasing rapidly and chromosome-scale, haplotype-resolved genomes are now within reach for many of these. This necessitates a transition from linear, single-reference to pangenome approaches in genomics. Graphs can represent such pangenomes, large numbers of related sequences, in a compact fashion. PanTools offers a practical pangenome sequence representation, indexed and stored in a graph database, annotated with structural and functional information.

In this work we have extended PanTools with read-mapping functionality. The method generates accurate alignments to all (or a subset) of the constituent genomes at once. Simultaneous mapping of reads allows avoiding redundant computations and can optionally distribute reads over genomes in a competitive manner, required in applications such as metagenomics. PanTools thus offers a solid basis, which can and will be further extended to integrate and mine different types of -omics data, paving the way towards comparative pangenomics.

## Methods

Before we present the read-mapping algorithm, first we briefly explain the pangenomic data structure to which the reads will be mapped. Then, we discuss our approach to competitive read mapping and finally introduce the data and methods used in the experiments.

### Structure of the pangenome

PanTools condenses multiple genomes in a generalized de Bruijn graph (gDBG), stored with structural annotations and proteomes in a graph database (24). There are two additional databases, an index database and a genome database, which facilitate efficient graph indexing and sequence retrieval respectively. A memory-mapped implementation of graph, index and genome databases minimizes the required I/O operations even in random access scenarios.

The gDBG captures the similarity and divergence of genomes at the resolution of *k*-mers. It is a compressed, bi-directed DBG, i.e. there is no non-branching path in the graph and every node represents a piece of double-stranded DNA of minimum length *k*, which occurs only once in the graph. Each sequence (contig, scaffold or chromosome) can be traversed as a continuous path in the graph in either forward or reverse direction. The positions of the node in the constituent sequences are stored on the edges of the graph. Figure 5 illustrates a node of this graph, a piece of DNA occurring in sequence 1 at position 8, sequence 2 at position 12, both in forward direction (TAC); and in sequence 3 at position 4, in reverse direction (GTA). During the gDBG construction we build a *k*-mer index that maps canonical *k*-mers to a unique graph coordinate: a triple of the identifier of the node, the zero-based offset of the *k*-mer in the node and the direction of the *k*-mer. For example, the 2-mer AC is mapped to coordinate (56, 1, F) if it occurs in node 56 at offset 1 in forward direction.

**Figure 5:**
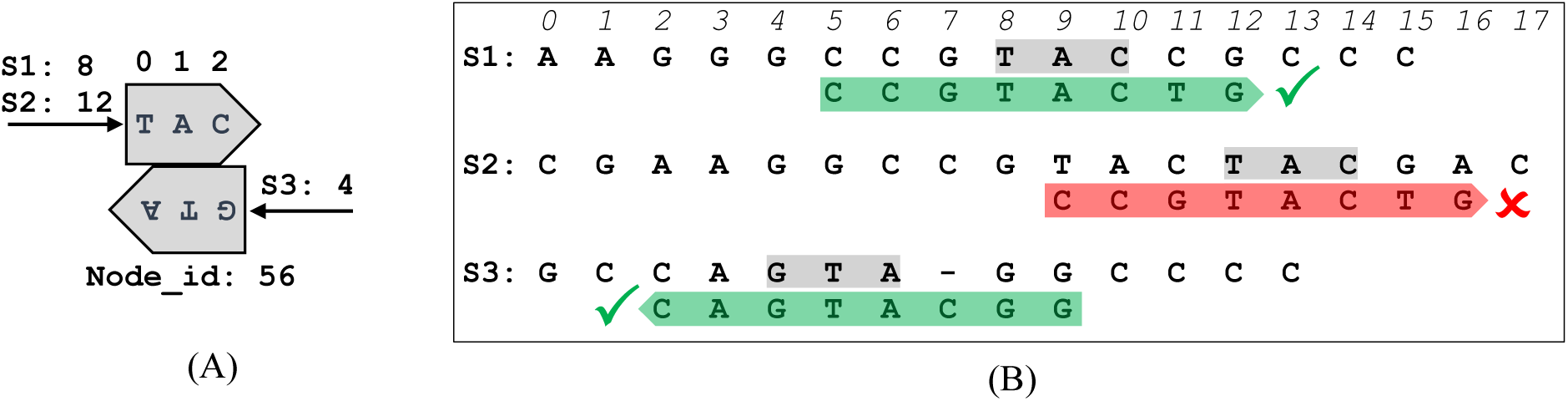
**A.** Structure of a node of generalized DBG; this piece of DNA occurs in three sequences S1, S2 and S3, respectfully, at positions 8, 12 and 4. **B.** After retrieving the candidate hits in each sequence and performing exact alignments, the read is mapped to S1 and S3 but not to S2.

### Read mapping

Given a genomic read, *α* (by default 15) equidistant *k*-mers are sampled from it and looked up in the *k*-mer index to retrieve the graph coordinates where the reads should map. The *k*-mer size used for read mapping is the same as the *k*-mer size used in graph construction, since the existing *k*-mer index is exploited. The number of *k*-mer samples *α* should be chosen higher when the error/mutation rate is high. The collected graph coordinates can then be translated into the start position of potential hits (the *candidate hits*) in the constituent sequences. For example, again consider 2-mer AC from read CCGTACTG. The position of AC in sequence 1 is the position of node 56 in this sequence (8) plus the forward offset of AC (1) in the node, i.e. 8 + 1 = 9. The offset of AC in the read is 4, so the position of the candidate hit on sequence 1 is 9 – 4 = 5. Candidate hits could be supported by different number of *k*-mer samples. Candidate hits are therefore sorted by the number of supporting *k*-mers (in decreasing order) and local alignments are only calculated for the first *ω* (by default 15) hits in this ordered list. Local alignment is performed using a Smith-Waterman algorithm with a banded matrix to reduce the number of calculations by limiting the number of gaps (by default 5) allowed in the alignment. As PanTools was developed for short reads, where a limited number of insertions/deletions is expected, this seems reasonable. If the alignment identity, defined as the number of identical positions divided by the length of alignment, is higher than a threshold *π* (by default 0.5), the hit will be reported as a *proper hit* (Figure 1B).

Algorithm 1 shows the pseudocode of our read mapping approach. Ideally, all *k*-mers, sampled from a read, point to the same position in a sequence. However, in the presence of sequencing errors, polymorphisms and genomic duplications some *k*-mers may not be found or may be found at multiple locations. Hence, these locations are clustered, collected in *Pos*[*S*], based on their proximity and the cluster size is considered as a score for that candidate hit. As many candidate hits may be false positives with low scores, only the *ω* most high-scoring hits are considered (Line 7). If all of these hits are supported only by one *k*-mer, potentially a low-complexity one, we just consider the first hit for the alignment. For each sequence, all proper hits, whose alignment identity is greater than a minimum threshold, are collected (Lines 8-9). Users have the option of reporting all the highest-scored hits (all-best), a random one if there are multiple best choices (random-best), or all the collected hits (all).

#### Algorithm 1: PanTools read-mapping pseudocode.

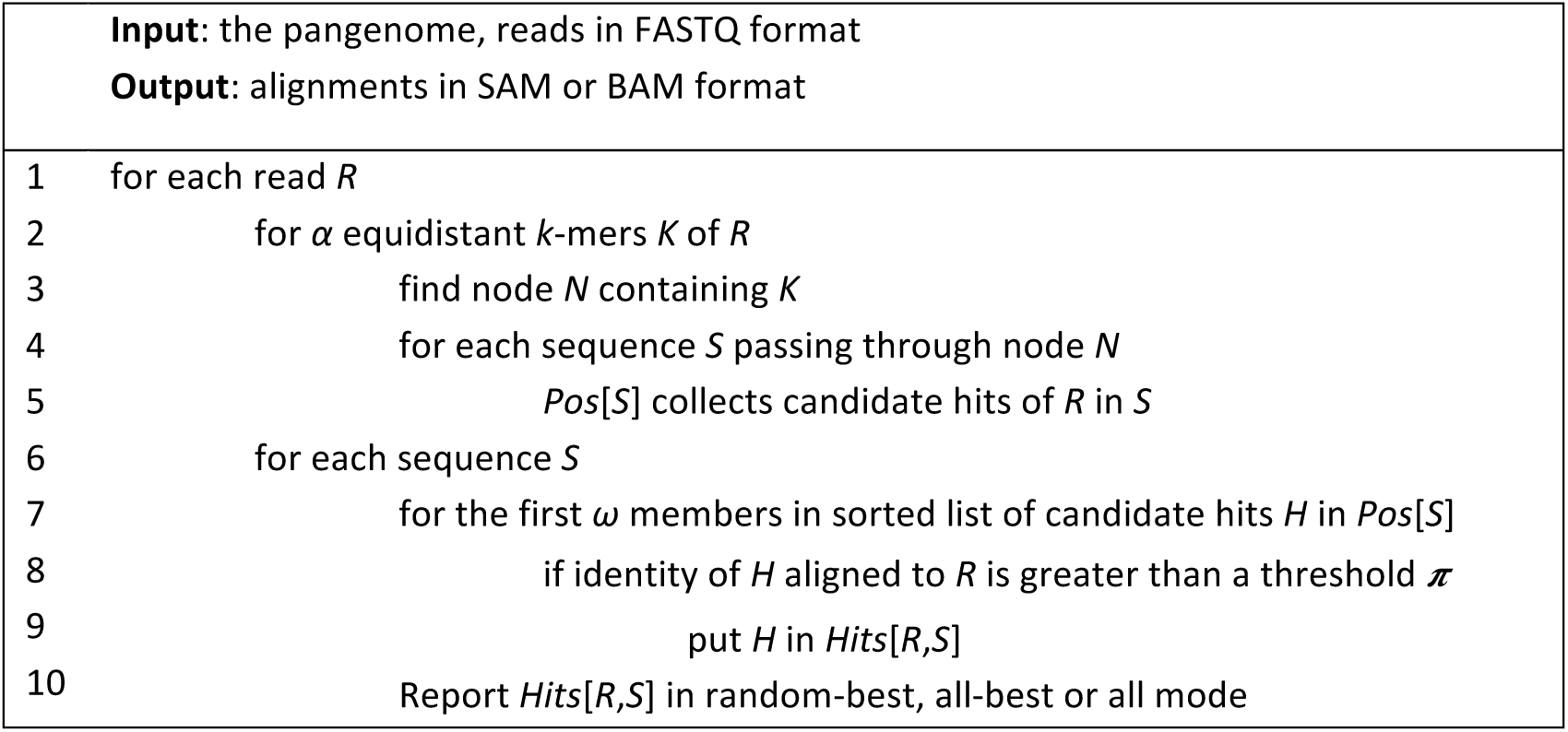

### Competitive mapping

PanTools is able to map reads in competitive mode, which is required for some applications, e.g. metagenomics and contamination screening. In this mode, proper hits to all the constituent genomes are collected and only those with the highest alignment identities (best hits) are considered. If there is a single best hit it is reported, otherwise PanTools offers three options for reporting the multiple best hits. First, *none-best* does not report any ambiguous hit; second, *random-best* selects one of the best hits randomly, either uniformly or based on some probabilities given to each genome; third, *all-best* which reports all the best hits. The random-best option works best for abundance estimation in metagenomics. When read mapping is followed by reference-guided assembly of the generated SAM files, it is preferable to use the all-best option to increase the horizontal coverage of the genomes.

### PanTools dependencies and parameters

For read mapping, PanTools depends only on KMC (25) for construction of the graph. The parameters affecting its mapping behaviour are listed in Table 5.

**Table 5:** Read-mapping parameters of PanTools.

| Parameter | Range | Default |
| --- | --- | --- |
| Number of parallel working threads | [1 .. cores] | 1 |
| Minimum acceptable identity of the alignment ($\pi$) | [0 .. 1) | 0.5 |
| Number of $k$-mers sampled from the read ($\alpha$) | (0 .. $r - k + 1$] | 15 |
| Minimum acceptable length of alignment after soft-clipping | [10 .. 100] | 13 |
| Maximum acceptable length of alignment | [50 .. 5000] | 1000 |
| Maximum acceptable length of fragment | [50 .. 5000] | 2000 |
| Maximum number of candidate hits to examine ($\omega$) | [1 .. 100] | 15 |
| Length of band in banded alignment | [1..100] | 5 |
| Stringency of soft-clipping | [0..3] | 1 |
| Alignment mode (Negatives for competitive, see manual) | [-3..3] | 2 |

### Data and experimental setup

All experiments were executed on an Ubuntu 14.04 server, Intel® Xeon® X5660@2.8GHz, with 6GB RAM and 16 processing cores. To generate synthetic reads, a 1% mutation rate was applied to the reference genomes of two model species *E. coli* (str. K-12 substr. MG1655) and *S. cerevisiae* (S288C, assembly R64) and 10x HiSeq 2500 (2×100) and MiSeq v3 (2×250) reads were simulated from the mutated genomes using the ART Illumina simulator (26). To show the accuracy and efficiency of PanTools, it was compared to four single-reference methods: Stampy, Bowtie2 and BWA-MEM and NextGenMap. The known genomic origin of the simulated reads allowed to compare the accuracy of the methods by counting the number of properly mapped (TP), wrongly mapped (FP) and unmapped (FN) reads, calculating the sensitivity = TP/(TP+FN) and specificity = TP/(TP+FP) of the tools which were then combined to an F-score = 2×sensitivity×specificity/(sensitivity+specificity) as the ultimate measure of accuracy.

To demonstrate the scalability of PanTools compared to the other tools, four sets of fungal genomes (pangenomes) were considered at different taxonomic levels. The first pangenome consisted of ten copies of the reference genome (R64) of *Saccharomyces cerevisiae* S288c. The second one contained ten different strains of *Saccharomyces cerevisiae* (including the reference R64). The third pangenome included genomes from ten different species in the *Saccharomyces* genus. Finally, the fourth pangenome contained genomes from ten different fungal genera in the family of Saccharomycetaceae (see Additional file 1: Experiment 2). For this experiment, the simulated MiSeq library of *S. cerevisiae* was used.

We demonstrated two real use cases on plant pangenomes. First, a recent llumina HiSeq 2500 paired-end sequencing archive (ERR2721960) of ∼13.3 million paired-end reads from DJA-1 accession was mapped to the reference and 18 additional high-quality assemblies of *Arabidopsis thaliana*. Second, three large paired-end sequencing libraries from the 150 Tomato Genome Resequencing Project (3) were mapped to the reference genome of *Solanum lycopersicum* (Heinz 1706) (27) and three additional accessions, *Solanum pennellii* (LA716) (28), *Solanum pimpinellifolium* (LA0480) (29), and *Solanum habrochaites* (LYC4) (3).

To test PanTools’ competitive mode of read mapping, a large library of 89.6 million paired-end reads from a stool sample was mapped (competitive random-best mode with uniform distribution) on a large pangenome of the reference genome database of the Human Microbiome Project. This database comprised of 130 archaeal strains over 97 species, 326 lower eukaryotes over 326 species, 3683 viral strains over 1420 species, and 1733 bacterial strains over 1253 species. The construction of this pangenome took 17 CPU hours, resulting in a database of size 104 GB, and read mapping was performed in 4.3 CPU hours. Additionally, we constructed a pangenome of the reference genomes from the “Critical Assessment of Metagenome Interpretation” (CAMI) benchmark, and compared our abundance estimates to those achieved by Kallisto quantification and DiTASiC on the three provided metagenomics datasets of low, medium and high complexity. Kallisto is based on a fast pseudo-alignment followed by an expectation–maximization (EM) approach to resolve the read abundance ambiguities. DiTASiC takes the raw pseudo-alignments of Kallisto, calculates the pairwise similarity of genomes and fits a generalized linear model (GLM) to resolve the read assignment ambiguities.

### Availability of data and materials

The read-mapping functionality has been implemented in PanTools version 2, available at https://git.wur.nl/bioinformatics/pantools, released under the GNU GPLv3 license. The datasets generated and/or analysed during the current study are available at: http://www.bioinformatics.nl/pangenomics.

## Funding

This work has been published as a part of the research project called *Pangenomics for crops* funded by the Graduate School Experimental Plant Sciences (EPS) in the Netherlands.

## Notes

https://git.wur.nl/bioinformatics/pantools

