## Supplementary figures and images for "Pangenomic read mapping"

### Supplementary Figure 1

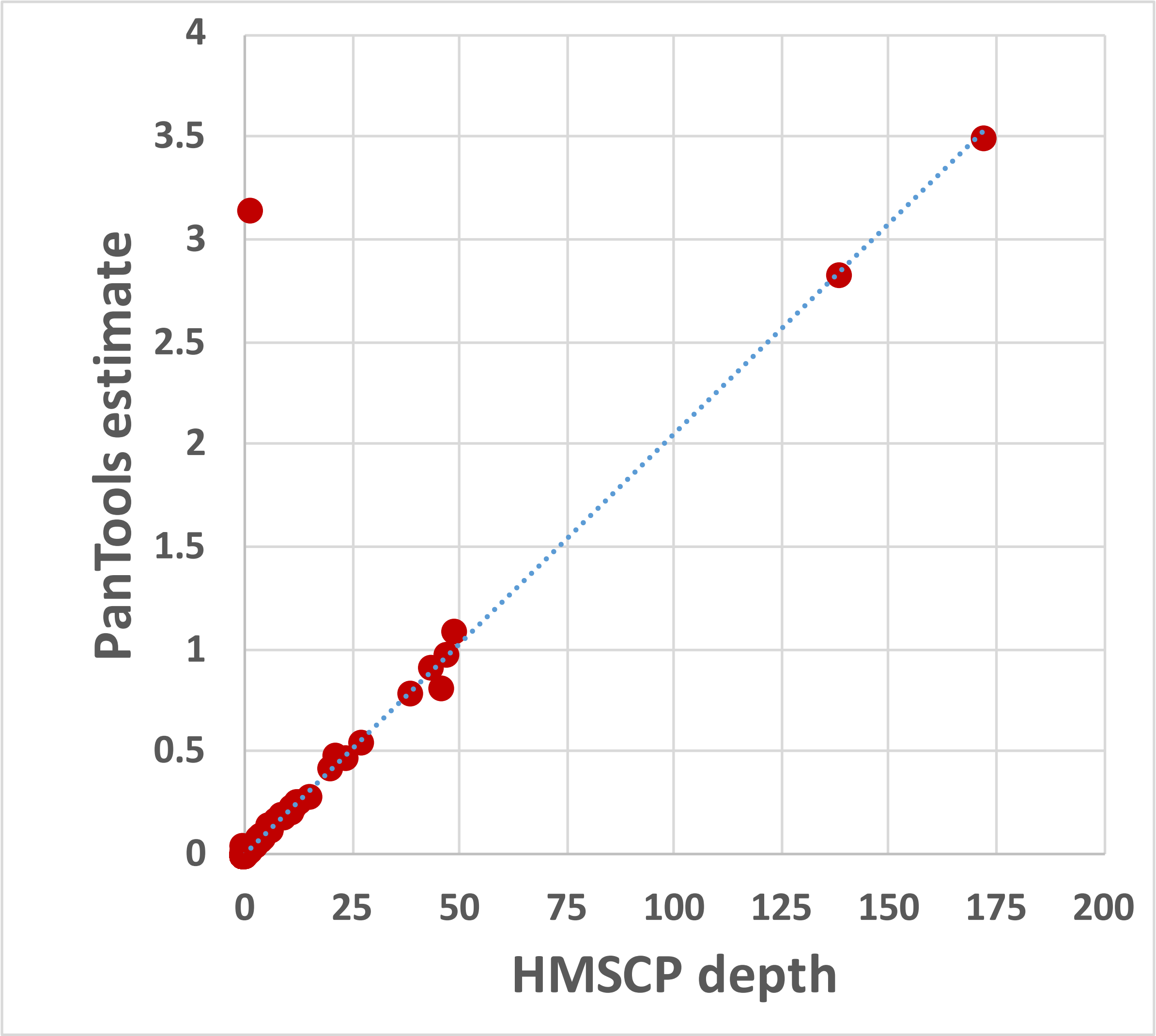


**Supplementary Figure 1:** Abundances as estimated by HMSCP and PanTools.
