## Supplementary Figure 2 for "Pangenomic read mapping"

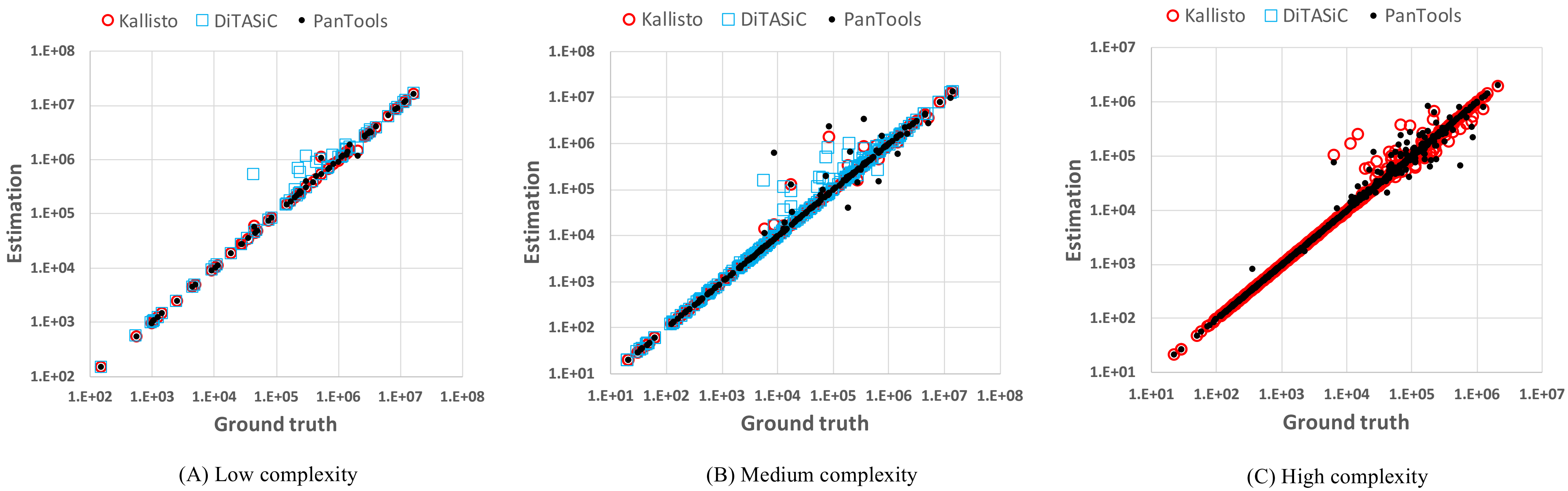


**Supplementary Figure 2:** Abundances as estimated by Kallisto, DiTASiC and PanTools versus the ground truth on the three CAMI benchmark data sets.
